# The interplay between multisensory integration and perceptual decision making

**DOI:** 10.1101/513630

**Authors:** Manuel R. Mercier, Celine Cappe

## Abstract

Facing perceptual uncertainty, the brain combines information from different senses to shape optimal decision making and to guide behavior. Despite overlapping neural networks underlying multisensory integration and perceptual decision making, the process chain of decision formation has been studied mostly in unimodal contexts and is thought to be supramodal. To reveal whether and how multisensory processing interplay with perceptual decision making, we devised a paradigm mimicking naturalistic situations where human participants were exposed to continuous cacophonous audiovisual inputs containing an unpredictable relevant signal cue in one or two modalities. Using multivariate pattern analysis on concurrently recorded EEG, we decoded the neural signatures of sensory encoding and decision formation stages. Generalization analyses across conditions and time revealed that multisensory signal cues were processed faster during both processing stages. We further established that acceleration of neural dynamics was directly linked to two distinct multisensory integration processes and associated with multisensory benefit. Our results, substantiated in both detection and categorization tasks, provide evidence that the brain integrates signals from different modalities at both the sensory encoding and the decision formation stages.

## Introduction

The different sensory channels provide complementary information, the integration of which leads to more accurate and faster behavioral decision (Welch and Warren, 1980; Stein and Meredith, 1993). The neural basis of multisensory integration and its loci in the hierarchy of brain computations have been the focus of myriad of studies (see for reviews (Talsma et al., 2010; ten Oever et al., 2016; Keil and Senkowski, 2018)). It is now well established that multisensory integration starts early in the process chain (Foxe and Schroeder, 2005; Schroeder and Foxe, 2005): both animal and human studies have demonstrated that the genesis of multisensory integration relies on cross-modal inputs to sensory cortices which informs about the spatiotemporal co-occurrence of sensory cues (Bizley et al., 2007; Lakatos et al., 2007; Kayser et al., 2008; Cappe et al., 2010; Mercier et al., 2013, 2015; Atilgan et al., 2018). At a later stage, the integration of information from different modalities pertains to congruency and reliability of multisensory inputs, as well as task relevance (Rohe and Noppeney, 2015, 2016; Kayser et al., 2017). However, it is unclear how multisensory integration interplays with decision making: it is still an open question whether the observed behavioral benefits of multisensory stimuli reflects a cumulative effect of multisensory integration at both sensory encoding and decision formation stages, and whether multisensory integration is at play during decision formation at all (Bizley et al., 2016).

Formation of perceptual decisions relies on the encoding of sensory evidence, which begins in the sensory cortices (Tsunada et al., 2016), is followed by interaction with parietal and frontal regions (Donner et al., 2009; Mostert et al., 2016), and then translated into behavior (see for reviews (Gold and Shadlen, 2007; Heekeren et al., 2008; Romo and de Lafuente, 2013)). This process chain of decision formation has been studied mostly in unimodal contexts and is thought to be supramodal (O’Connell et al., 2012; Romo and de Lafuente, 2013). Critically, the wide networks underlying perceptual decision making overlap with the neural substrates of multisensory integration (Ghazanfar and Schroeder, 2006; Romo and de Lafuente, 2013; Bizley et al., 2016), yet it remains to be established how the two processes are linked. This question is best exemplified by the longstanding debate about how the brain operates decision formation in a multisensory context: whether two decision formation processes operate independently for the different modalities as assumed by the parallel race model (Raab, 1962) or whether evidence is combined across modalities before feeding one decision formation process as in the co-activation model (Miller, 1982).

Given these two processing stages (sensory encoding and decision formation), the effect of multisensory integration on perceptual decision making could take place (Figure 1A): (1) during sensory encoding only that is before a supramodal decision formation, (2) during a multimodal decision formation, or (3) both during sensory encoding and decision formation. To test these alternative hypotheses, we employed a time-resolved decoding approach on human EEG while subjects detected or categorized unpredictable unisensory-cue (auditory / visual) or multisensory-cues embedded within a stream of audiovisual noise. Using multi-variate pattern analysis (MVPA), we first characterized sensory encoding and decision formation processes when only unisensory-cue was available in the noise (unisensory classifier). Thereafter, we applied the unisensory classifier onto multisensory-cues condition. This cross-condition decoding generalized in time revealed an acceleration of both sensory encoding and decision formation for multisensory-cues condition as compared to unisensory-cue condition. Lastly, we performed a direct decoding procedure between unisensory trials and multisensory trials and revealed two periods of multisensory integration intimately linked to sensory encoding and decision formation. Together, these findings, verified in both detection and categorization tasks, demonstrate that early multisensory integration speeds-up sensory encoding and therefore leads to an earlier onset of decision formation, while later multisensory integration speed up the rate of decision formation.

**Figure 1.**
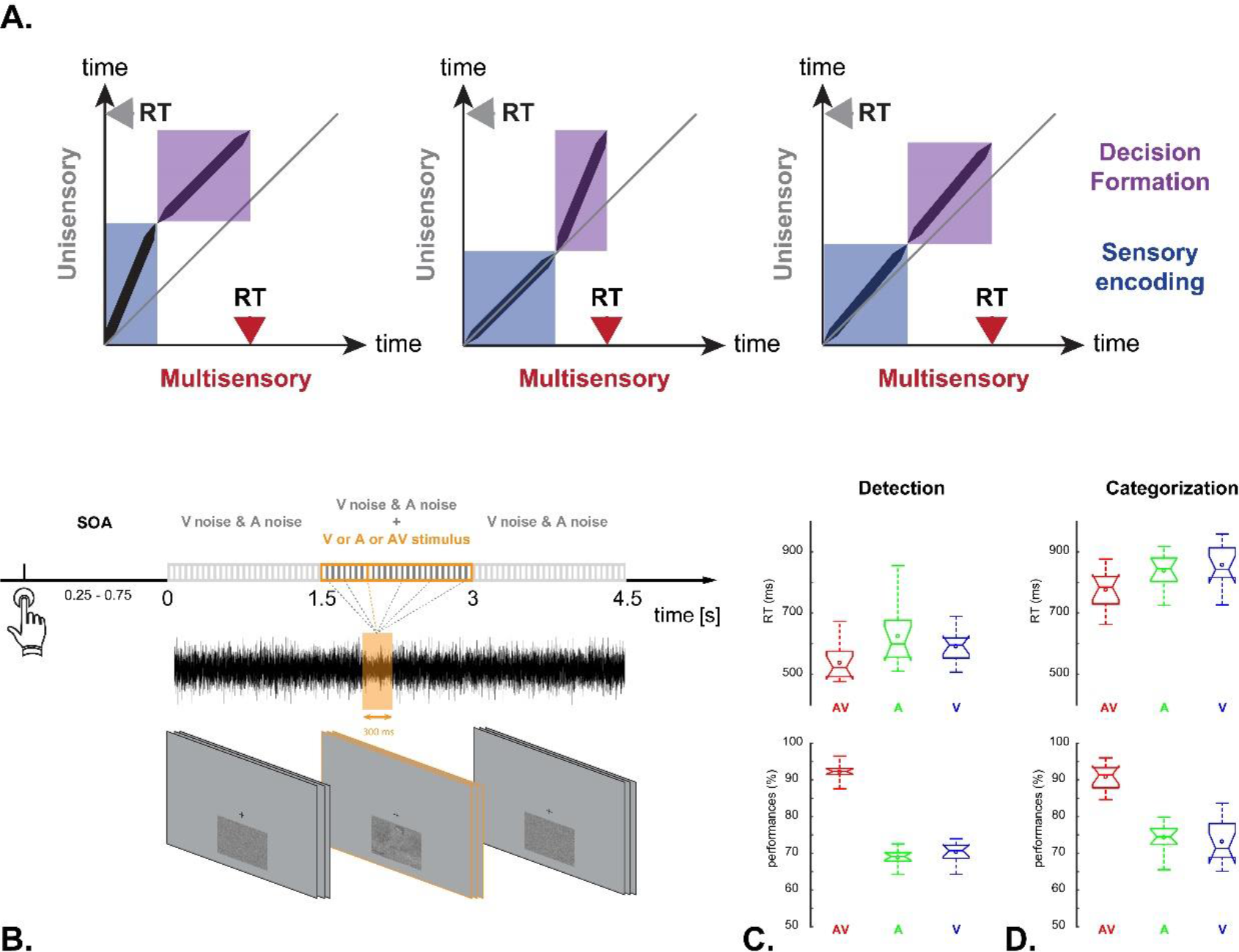
(A) Hypothesized effect of unisensory and multisensory signals on perceptual decision making dynamic during sensory encoding (in blue) and decision formation (in purple). Left: Once received, sensory inputs are fully integrated to reduce uncertainty and accelerate sensory processing, which in turn fuels a supramodal decision process. Middle: Information from the different senses is processed in parallel during sensory encoding and facilitatory effect of multisensory integration takes place only during decision formation stage. Right: The different modalities are combined at both processing stages. (B) Experimental design: participants had to detect or categorize an unpredictable cue fading in/off from a continuous stream of audio-visual noise. (C) Behavioral performance: Response time and accuracy distributions plotted for audiovisual (red, AV), auditory (green, A) and visual (blue,v) cue conditions during detection and categorization tasks.

## Material and Method

### Participants

Data were collected from 12 subjects (4 females, mean age 26.7 years) in the main EEG experiments. All had normal audition and normal or corrected-to-normal vision. The study was conducted in accordance with the Declaration of Helsinki and approved by the Inserm (Institut National de la Santé et de la Recherche médicale) ethical committee (Institutional Review Board IRB00003888 - agreement n°14-156). Written informed consent was obtained from all volunteers before the experiment.

### Experimental setup

EEG was acquired from 128 electrodes and 4 additional electrodes places above and below the dominant eye were used to record EOG. Signals were digitized at 2048 Hz, 24-bit A/D conversion (BioSemi ActiveTwo system, Netherlands). During the experiment, participants were seated comfortably in a quiet dark room. A chin-rest was used to maintain correct head position while fixating on a black cross continuously displayed on a gray background (BenQ XL2411, refresh rate: 100 Hz, resolution: 1920×1080). Instructions and visual stimuli were presented on a screen located 80 cm from the participant. Sounds were delivered through earphones (Etymotic ER.4), using a dedicated audio card (Sound Blaster Audigy 5/Rx). Responses were collected through a numeric keypad. The experiment was programmed and controlled using Presentation software (version 18.xx from NeuroBehavioral Systems, USA).

### Stimuli

Stimuli were primarily from the CerCo databases (the Brain and Cognition research Center, CerCo, UMR 5549 CNRS). They were typical sounds and images of living or inanimate categories (respectively birds, dogs, monkeys and cars, guitars, phones). All stimuli were equated using the following procedure. Sounds were first calibrated (11025Hz, 16 bits, mono) and then rms-normalized. Auditory noise was added by permuting samples with a morphed average of all sounds. Images were first cropped (400×300 pixels) and converted to black and white. Intensity range was normalized and image histograms equalized to the mean image. Visual noise was created by shuffling pixels. For both sounds and images, the signal-to-noise ratio (SNR) ranged from 0% to 100% in 5% steps increment.

The stimuli used for the main experiment were selected from a larger pool of stimuli after piloting on a separate group of participants (n=8). Every selected item reached at least 50% of correct recognition at 50% of SNR in the pretests to insure representativeness and homogeneity of the final pool. Following the piloting, the selected set of stimuli contained 60 sounds and 240 images with an equal number of items per category.

During the main experiment, unisensory performance accuracy was kept at 70% through a continuous staircase procedure (adaptive up-down method (Levitt, 1971) computed separately for each category in unisensory conditions).

### Experimental design

At the beginning of each trial the participant was prompted to press a button. Following a random SOA (250-750ms), an audio-visual sequence started. Each sequence lasted 4.5 sec and contained dynamic audio-visual noise. At any random time between 1.5 sec and 3 sec, an unpredictable signal cue stimulus was presented in any modality (auditory, visual or both at the same time) on 85% of trials. The remaining 15% were trials with audio-visual noise only (catch trials).

During the audio-visual sequence, smooth transition between noise and signal cue was handled through a gradual increase/decrease of signal-to-noise in the course of target presentation (300ms cycle for signal fading-in/off). The visual stream was constructed by displaying at every screen refresh a random picture with a given signal-to-noise. For each sequence, the pictures used where built from one of the selected images (with 500 versions at 0% signal-to-noise and 100 versions for each signal-to-noise above 0%). Each auditory stream was constructed by concatenating the different noise version of a selected sound (either at 0% signal-to-noise or at a given signal-to-noise for signal cue).

Every participant performed two tasks. In the detection task, participants had to indicate whether a trial contained a signal cue or not in either or both modalities by pressing a response button. In the categorization task, participants had to indicate if the signal cue was an animal or an inanimate object by pressing the corresponding response button (counterbalanced across participants). In case of audio-visual cues, the two modalities were congruent (*e.g.*, the image of a bird was presented with the sound of a bird). Responses given at the end of a stimulus sequence were not taken into account. Task order was counterbalanced between participants. Each task was divided into five blocks, each containing about 160 sequences. To maintain vigilance, participants were encouraged to make self-paced breaks after each block.

To assess multisensory effect at the behavioral level, both accuracy and responses times from the multisensory-cues condition were compared to the unisensory-cue conditions using paired random permutation test (10000 iterations).

### Signal preprocessing

Electrophysiological data (EEG, EOG) were scanned using a semi-automatic artifact detection procedure based on signal statistics (variance, amplitude and z-score computed across trials and channels separately). To exclude remaining artifacts, continuous data were visually inspected. Artifacted channels were interpolated using spline method. Then average reference was applied. Ambient noise (50 Hz and 100Hz-150Hz harmonics) was removed by fitting a sine and cosine at the specified frequency to the single trial data and subsequently subtracting the estimated components. Trials were defined from +/−1.625 from stimulus onset fade-in. For “catch” trials (i.e. only audio-visual noise, no target), a sham onset time point was randomly picked between the 1.5-3s period. Trials were demeaned and high-pass filtered at 0.5Hz using a windowed Sinc FIR filter. Trials with extreme target stimulus signal-to-noise ratio were discarded on the basis of two signal-to-noise steps above and below the median in each category (in either the visual or the auditory condition). The data were processed offline by using custom-written scripts in MATLAB (MathWorks, MA, USA), the FieldTrip Toolbox (Oostenveld et al., 2011) and the LIBLINEAR library for large linear classification (http://www.csie.ntu.edu.tw/~cjlin/liblinear).

### Multivariate pattern analysis

#### Principle

The aim of the MVPA approach is to best exploit the information provided by every sensor to isolate activations that are specific to a given brain operation. When applying MVPA to EEG, the time series from all electrodes are optimally combined to define topographical weights that maximally discriminate experimental conditions at a given time point. In the present study, to obtain these topographical weights, we used a linear classifier based on L2 regularized logistic regression (Fan et al., 2008) in a *Monte-Carlo* stratified cross-validation procedure (hold-out, 200 iterations).

#### Procedure

Data were down-sampled (128 Hz) to reduce computation time while maintaining sufficient temporal resolution (Grootswagers et al., 2017). Trials were defined either relative to the cue fade-in onset (−100 to 1300 msec) or relative to the response (−1500 to 200 msec). Each trial was baselined using −100-0 msec interval (cue-locked analysis) or the entire epoch (RT-locked analysis). On each cross-validation iteration (CV), the data-set was randomly split into a training set (90% of the trials) and a testing set (the remaining 10% of the trials), each condition being equally represented by the same amount of trials (stratified cross-validation). Last, the signal at each electrode was normalized across trials using the estimates from the training set (Crouzet et al., 2015; Edwards et al., 2018).

#### Weights projection

For each time point, a weight was assigned to each electrode which reflects how this feature contributed to maximizing the decoding. Topographical weights were transformed back into activation patterns by multiplying them with the covariance in the data and applied to the single-trial time series to track the temporal course of the cognitive operation isolated by the corresponding classifier (see below).

#### Time generalization

Classifiers that best differentiate conditions at a given time point were tested on every other time point, leading to a “temporal generalization” matrix. This step was performed within the cross-validation iteration which implies that trials used for training and testing (at the same/different time point) were from different trial sets. Such temporal generalization permits to draw the blueprint of brain processes by distinguishing canonical motifs that are not accessible otherwise. Similar variation in decoding performance over time can originate from different scenarios revealed through temporal generalization. For instance a chain of different processes can have the same profile of decoding performance as the reactivation of a unique process. In the former case temporal generalization would show a diagonal motif, in the second case a checkerboard like motif where the process is repeated (King and Dehaene, 2014).

#### Generalization across conditions

To assess brain responses similarity and/or specifics across conditions, the classifiers obtained when training in one condition (*e.g.*, response time decoding of unisensory-cue trials) were tested for generalization to another conditions (not being part of the testing set or not being directly tested, *e.g.*, response time decoding of multisensory-cues trials). This step was performed at each iteration on normalized data. We also tested for the temporal generalization across conditions, following the same procedure as for within-condition temporal generalization described above. Generalization across time and experimental conditions capture the neural architecture of brain operations and reveal how processing stages are modulated between experimental conditions (Marti et al., 2015; Myers et al., 2015; Salti et al., 2015; King et al., 2016).

#### Statistical validation

For each iteration the classifier generated a measure of the classification accuracy. That is the proportion of test trials that were correctly classified using the topographical weights computed from the training set. Chance levels were obtained by running the same procedure while shuffling the labels between conditions. As within-subjects analysis, classifier performances from each iteration were used to compute confidence intervals of the mean using a bootstrapping procedure (95% CI from 1000 boostraps) which was then compare with results from shuffled data to estimate significant decoding. Between-subject analysis was performed using a one-tail paired random permutation test on the mean performance across CV iterations for real and shuffled conditions (10000 iterations). To control for multiple comparison a cluster-based correction was applied with false alarm rate set at 0.005% cut-off (Maris and Oostenveld, 2007).

#### Representativeness of classification components

To explore the relevance of the topographical weights defined by the classifier, we applied the weights to single-trial time series. This approach leads to a one-dimensional projection of the classifier and allows to quantify its representativeness as a function of signal-to-noise-ratio (Figure 2 B-C, supplementary Figure 1 B-C). The same approach was used to assess the dynamics of decision formation as a function of response time and to compare unisensory and multisensory conditions. After projecting the weights of the classifiers based on response times regression, single-trial time courses were binned by RT (same amount of trials in each bin: fast, medium, slow), smoothed (using a gaussian-weighted moving average over 5 time points), baselined corrected (from −100ms to cue fading-onset) and averaged. A linear regression was computed between the onset of the rising component and its maximum absolute amplitude. Onset was defined by an absolute amplitude larger than the mean baseline amplitude plus-minus two standard deviations. Finally, the slope was compared between RT bins (fast, medium and slow) and conditions (unisensory or multisensory) using a two-way analysis of variance with participants as repeated measure.

**Figure 2:**
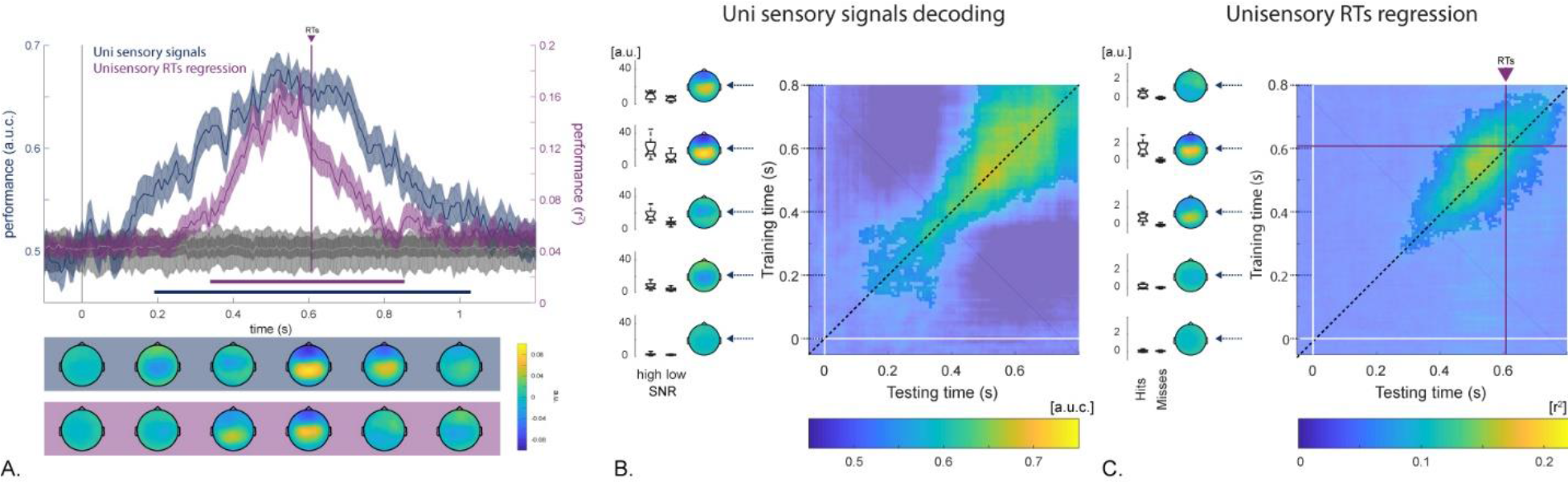
Detection task: time-resolved pattern classification of unisensory signal cue encoding and decision formation. (A) Mean decoding performance as a function of time (+/− 95% mean confidence intervals). In blue: decoding of unisensory signal cue trials against noise trials. In purple: decoding based on response times. For both, chance levels are depicted in gray. Vertical line represents subject-average response time. Color-coded lines below the x-axis signify statistically significant decoding performance compared to the chance level (corrected for multiple comparisons across time, p < .005). Topographical plots underneath depict activation patterns corresponding to the classifier weights (mean over 200 ms time window). Temporal generalization matrix and corresponding activation patterns depicted for the classifier decoding unisensory trials from trials containing audio-visual noise only (A), and for the classifier decoding response times. Box plots illustrate the consistency of activation patterns across single trials binned by the amount of signal to noise ratio of the cue or between single trials hit or miss.

#### Link between classifiers

For each subject, classifiers showing the highest decoding performance during the sensory encoding stage (0 to 300 ms) and the decision formation stage (300ms to the response) were identified. Then the corresponding weights were projected onto multisensory-cues trials (hits only) to obtain a single-trial estimate of each classifier (*i.e.*, decoding sensory encoding, decision formation and early/late multisensory integration). The maximum absolute amplitude of the one-dimensional decoder projection was extracted for each single trial and compared using a correlation analysis. Given the hypothesizing positive relationship, one-tail Pearson correlation was computed. Single-trial level analysis was verified at the population-level using the mean across trials.

## Results

### Behavior

We applied decoding analysis to EEG signal recorded while participants had to detect (task 1), or to categorize (task 2), a target-cue embedded in a stream of audio-visual noise (Figure 1B). The signal cue was either visual, auditory, or audio-visual, and could be presented at any moment in the stream of noise or be absent (catch trials). Signal cue consisted of an unpredictable faint gradual increase-decrease of signal-to-noise-ratio (SNR). The amount of SNR was titrated to maintain performance at an accuracy level of 70% in either unisensory conditions.

Behavioral analysis revealed typical multisensory benefit effect (Figure 1C), with greater accuracy following multisensory cues in both detection (p-values < 0.001) and categorization tasks (p-values < 0.001). Mean response times were faster in the multisensory condition as compared to both unisensory conditions (detection task: p-values < 0.001; categorization task: p-values < 0.002). These findings demonstrate that both detection and categorization of the signal cue embedded in the audio-visual noise is facilitated and fastened when cues originate from two modalities simultaneously.

### EEG decoding analysis of sensory encoding and decision formation following unisensory signal cue

To characterize the functional processing stages at play following unisensory cue we used a series of multivariate pattern analysis (MVPA). Firstly, a linear multivariate classifier was trained to distinguish trials containing unisensory signal cues (auditory / visual) embedded in audio-visual noise from trials containing only audio-visual noise. Time resolved decoding performance gradually increased above chance level following unisensory cue onset, peaked and returned to chance level (see Figure 2). Secondly, we performed a non-binary classification to decode correct unisensory trials as a function of response times (RT) and thus evaluate the formation of decision over time. Compared to the first classifier, classification performance of this RT-based classifier rose later above chance but peaked at the same latency before the behavioral response was made (Figure 2A and supplementary Figure 1). To facilitate the comparison between the results from the two classifications and further delineate the temporal extent of brain processes related to perceptual decision making, we computed the topographical representation of classifier weights (activation patterns) and performed temporal generalization. Temporal generalization matrix is obtained by testing across all time points a decoder trained at a given time point and thereby characterizes canonical motif of neural operation (e.g. sustained, chained or reactivated see (King and Dehaene, 2014) for a comprehensive review). The activation patterns and the temporal generalization matrix captured by the unisensory signal cue vs. noise classifier revealed two processing stages. The first one consisted in a prominent parietal negativity, similar to the early target selection signal, referred as the N2 component (Luck and Hillyard, 1994; Gamble and Woldorff, 2015; Loughnane et al., 2016). The second stage consisted in a positive centroparietal topography similar to the centroparietal positivity component (CPP) recently described as a hallmark of decision formation (O’Connell et al., 2012; Twomey et al., 2016). This second stage was also isolated by the RT-based classifier confirming its role in the decision process preceding the behavioral response (Figure 2 B-C and Supplementary Figure 1). Last, as evaluated by projecting decoders’ weights on high and low SNR single trials, both stages appeared to be modulated in strength by the amount of sensory information (see box plots in Figure 2 B-C and Supplementary Figure 2 B-C) as reported for the N2 and the CPP components (O’Connell et al., 2012; Loughnane et al., 2016).

### Decoding generalization to multisensory signal cues reveals acceleration at both sensory and decision stages

Next we sought to uncover the neural mechanisms underlying behavioral performance benefit following multisensory cues by applying a cross-condition decoding approach. Classifiers derived from unisensory trials were used to discriminate trials containing multisensory signal cues versus trials containing only audio-visual noise. Moreover, to account for possible temporal differences across conditions and thus accommodate the decoding of brain operations happening at different latencies we further tested the capacity of classifiers to generalize across time. Generalization across time and experimental conditions captures the neural architecture of brain operations and reveal how processing stages are modulated between experimental conditions (see instantiation in: (Marti et al., 2015; Myers et al., 2015; Salti et al., 2015; King et al., 2016)). This approach revealed that classifiers trained on unisensory trials were able to decode multisensory trials effectively. This indicates that unisensory and multisensory decision making follows a similar time course. Furthermore, matrix generalization unveiled an off-diagonal pattern: unisensory classifiers led to higher decoding performance at earlier latencies for multisensory trials (Figure 3 and Supplementary Figure 2). As evidenced by classification against audio-visual noise (Figure 3A) and RT-based decoding (Figure 3B), this acceleration pattern occurred during both sensory encoding and decision formation stages. In order to verify that the speed-up significantly increased with time we calculated the distance between the two decoding time courses every 100ms (*i.e.*, orange area in Figure 3 and supplementary Figure 2). This integral between significant decoding performances was found to increase linearly with time (Pearson correlation: detection task: p = 0.002, rho = 0.91; categorization task: p = 0.01, rho = 0.83). Thus, the present results demonstrate that combination of multisensory-cues speeds-up neural dynamic all along the course of processing and thereby strongly supports the view that multisensory integration processes are at play during sensory encoding and during decision formation (third hypothesis depicted in Figure 1A).

**Figure 3:**
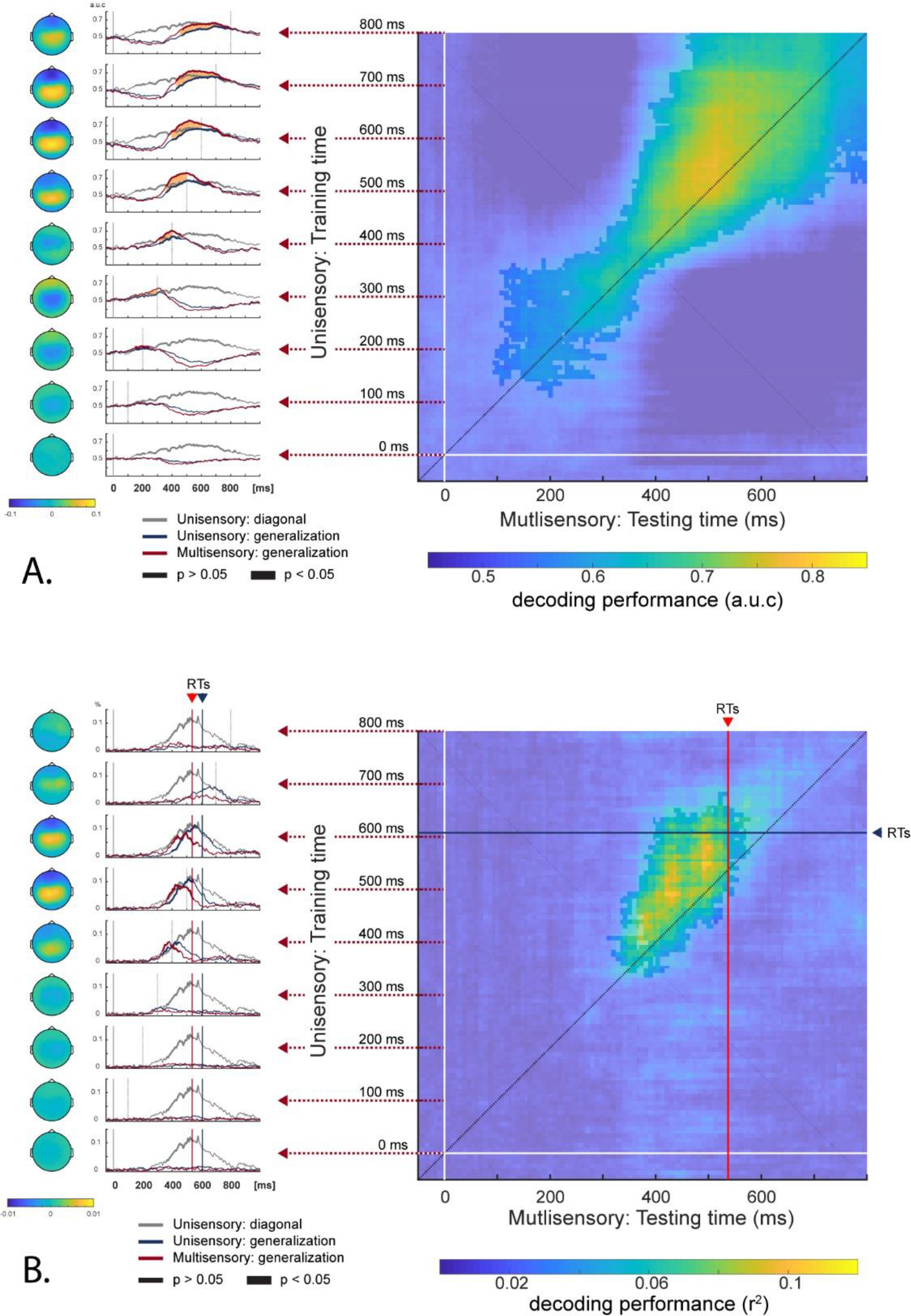
Detection task, generalization from unisensory to multisensory condition. Classifiers trained on unisensory trials were tested (A) to distinguish multisensory signal cue trials vs. trials containing audio-visual noise only, or (B) to decode response times of multisensory-cue trials. Right side: Classifiers trained at a given time (y-axis) are tested at every other time moment (x-axis). Blue horizontal and red vertical lines represent mean response times for unisensory trials and multisensory trials respectively. Left side: Each row represents activation patterns and decoding performance of classifiers tested on multisensory trials (in red) and on unisensory trials (in blue), plotted every 100 ms of the training time (with 0 ms on the bottom). The grey lines represent decoding performance for unisensory trials along the diagonal of the temporal generalization matrix (dotted line along the diagonal in Figure 2). Orange areas indicate periods of process acceleration.

To further investigate the results of the RT-based classification, we projected the weights of the RTs-based classifier on single trials. The obtained one-dimensional projections of the decisional process were then binned by RTs (fast, medium and slow quantiles) and subjected to a linear regression to access the slope of decision formation. These slopes were estimated separately for unisensory and multisensory trial hits and then subjected to a two-way analysis of variance with subjects as a repeated measure. Results showed a significant effect of RT-binning on slopes: decision formation time-courses were steeper for faster trials and less steep for slower trials (detection task: F = 3.6, p = 0.0336; categorization task: F = 5.9, p = 0.00442). Moreover, results revealed an effect of signal cue on the slope: decision formation buildup more rapidly in multisensory-cues trials than in unisensory-cue trials (detection task: F = 8.3, p = 0.00535; categorization task: F = 7.8, p = 0.00688). This last analysis illustrated that response time varies with the rate of decision formation and uncovered that the rate of decision formation is faster for multisensory-cues.

Temporal generalization across conditions illustrates the chain of processes common to unisensory and multisensory conditions. However, neural activity specific to multisensory processing (*i.e.*, multisensory integration) was not targeted using this approach. To examine if the acceleration of brain network activation in multisensory-cues condition is the only difference with unisensory-cue condition, we compared the temporal generalization across conditions with direct decoding of multisensory-cues condition. That is, we trained classifiers to distinguish multisensory signal cues trials from audio-visual noise only (multisensory classifiers). Thereafter, we compared decoding performance of multisensory classifiers with the highest off-diagonal performance from the generalization matrix across time and condition (see Figure 3). Comparison of indirect and direct decoding showed a similar time-course (Supplementary Figure 3). However, during two time periods decoding performance of multisensory classifiers (direct approach) was significantly higher than that obtained from temporal generalization across condition (indirect approach); even though temporal differences between conditions were accounted for. This result indicates that decoders built from multisensory trials extracted neural activity which could not be captured by the decoders build from unisensory trials alone, neural activity likely related to the integration of multisensory-cues (i.e., related to multisensory integration processes).

### Time-resolved decoding of multisensory integration

To tackle multisensory integration, we trained classifiers to differentiate between correct multisensory-cues trials and correct unisensory-cue trials. Time-resolved pattern classification showed above chance performance starting early after the cue fade-in onset, followed by two peaks and characterized by two activation patterns. Early period presented a wide parieto-occipital negativity while the late period showed a centro-parietal positivity, with both topography being asymmetrical with larger activity on the right side (see Figure 4A and supplementary Figure 4A). To assess if the late period, showing the highest decoding performance, was caused by a time lag between conditions (due to the speed-up reveal by the temporal generalization across conditions, see above), we ran the same analysis using trials time-locked to the response. This second analysis confirmed the existence of two periods where multisensory integration occurs, each characterized by a peak in the decoding performance and substantiated by a specific activation pattern. As a control analysis, we tried to decode missed multisensory-cues trials versus missed unisensory-cue trials and did not find above chance performance. Such null result sets a parallel between classifier efficiency in decoding multisensory signal and the possible absence of multisensory integration when signal cues were integrated effectively. This inference is further supported by the link between classifiers decoding performances and multisensory gain on behavioral performance. That is, we found a positive correlation (r^2^=0.62, p=0.015) between classifier performance and the difference in behavioral performance between multisensory and unisensory conditions (*i.e.*, multisensory gain).

**Figure 4:**
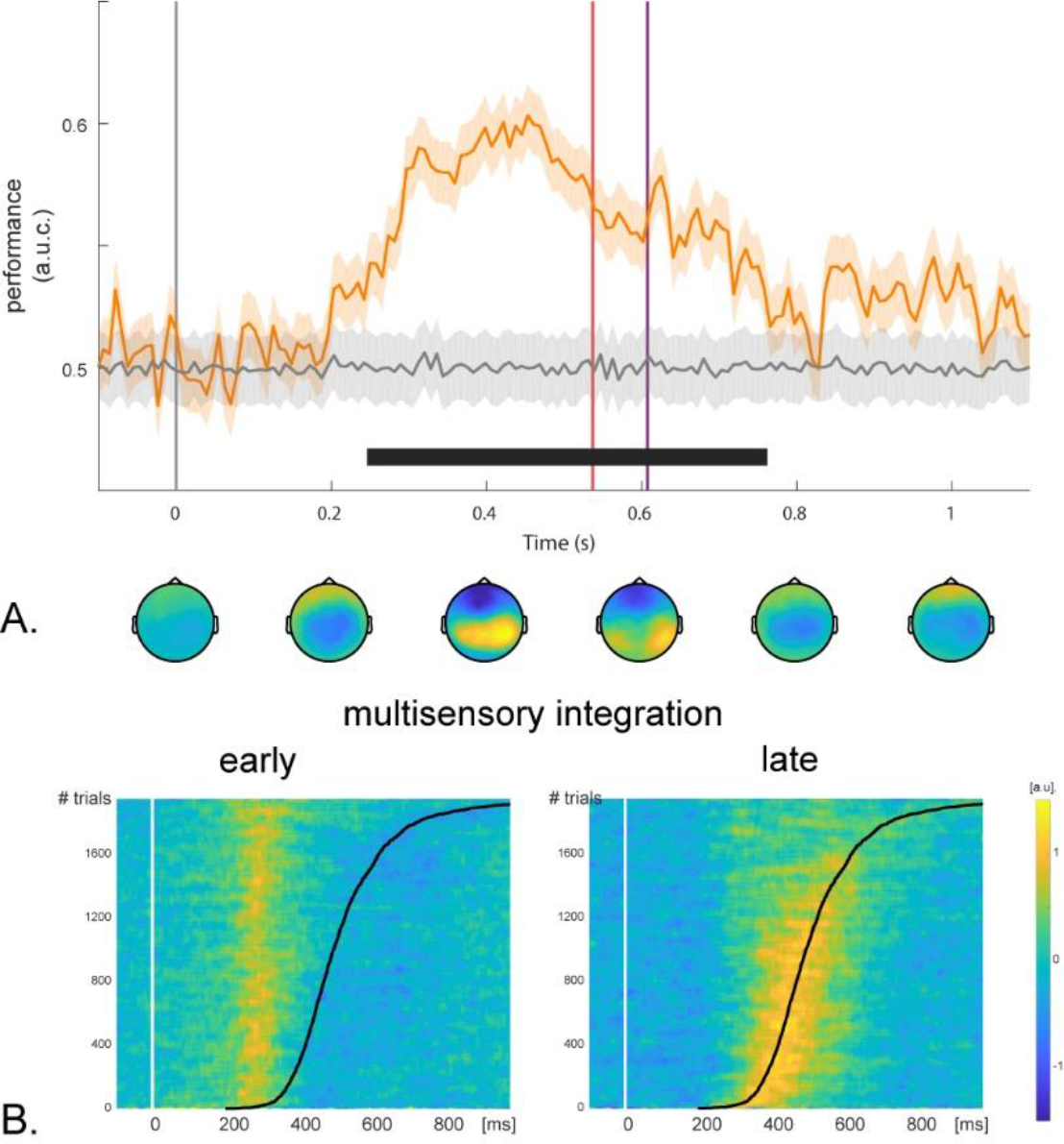
Detection task: time resolved decoding of multisensory integration. (A) Mean classification performance (orange line) as a function of time (+/− CI mean; chance levels in gray.), with the black line under the x-axis indicating statistically significant decoding performance compared to the chance level (corrected for multiple comparisons across time, p < .005). Topographical maps underneath depict the activation patterns from time-resolved classification procedure (mean over 200 ms time window). Vertical lines represent mean response times for multisensory trials (in red), and unisensory trials (in purple). (B) Single-trial projection of channel weights from classifiers decoding early or late multisensory integration periods. Single trials are sorted by response times (black line).

### Multisensory integration linked to sensory and decision stages

After characterizing the two activation patterns related to multisensory integration (Figure 3), we assessed how they were associated with the identified processing stages of perceptual decision making: sensory encoding and decision formation stages. For each participant, we identified the classifier weights leading to the highest decoding performance during periods of sensory encoding and decision formation (Figure 3 and 4) and applied these weights to the multisensory-cues trials. This procedure provided us with single-trial estimates of each classifier over time, which in turn allowed us to evaluate the relationship between the corresponding cognitive processes (early/late multisensory integration and sensory encoding/decision formation). That is if the signal strength of sensory encoding and the signal strength of decision formation were linked to the efficiency of multisensory integration. This comparison revealed a significant linear relationship between early multisensory integration and sensory encoding (detection task: r=0.5273, p=0.0391; categorization task: r=0.7920, p=0.0011), and late multisensory integration and decision formation (detection task: r=0.7920, p=0.0011; categorization task: r=0.5015, p=0.0483). Lastly, to evaluate the influence of multisensory integration on RT over time, we sorted single-trial time series of the corresponding classifiers by RT (Figure 4B). This visualization revealed that the second period of multisensory integration accounted for trial-to-trial RT variability. Overall these results revealed a link between the cognitive processes isolated by the different classifiers: while early multisensory integration was related to sensory encoding process, later multisensory integration was linked to decision formation and to response time variations.

## Discussion

Multisensory-cues lead to faster and more accurate behavioral performance as compared to unisensory-cue. However, the origin of such multisensory benefit is still unclear and could arise from integration happening during sensory encoding and/or during decision formation. In the present study we aimed to pinpoint the critical time periods during which multisensory integration influences perceptual decision making. Using multivariate pattern analysis (MVPA) we first characterized the functional stages of unisensory perceptual decision making: sensory encoding and decision formation, we then demonstrate that multisensory benefit originates from the acceleration of both processing stages following the presentation of multisensory-cues, finally we uncovered that processing stage acceleration is tightly linked to distinct multisensory integration processes at play during sensory encoding and decision formation.

### The course of perceptive decision making following unisensory-cue

Pioneering single-unit animal studies revealed that decision-related activity recorded in lateral intraparietal cortex and frontal eye fields paralleled temporal accumulation of sensory evidence leading to the behavioral response (Gold and Shadlen, 2007; Heekeren et al., 2008). Neural correlates of perceptual decision making have since been identified in multiple brain areas, with neural code of decision making process spanning across cortical hierarchy - from sensory areas to parietal and frontal regions (de Lafuente and Romo, 2006; Siegel et al., 2015). In humans, a series of seminal EEG studies identified a brain signal, the centro parietal positivity (CPP), presenting many characteristics of a neural signature of decision formation (O’Connell et al., 2012; Twomey et al., 2016). In our study, the activation pattern and the time-course of decoding performance of the RT-based classifier (Figure 2 and supplementary Figure 1) match the topography and temporal characteristics of the CPP: centro-parietal topography and progressive increase in amplitude, peaking before the behavioral responses. Moreover, similar to our finding (see Figure 2 and supplement Figure 2), the CPP is modulated by the amount of sensory evidence available in the stimulus and differentiates hits from misses (O’Connell et al., 2012; Twomey et al., 2016). These analogies further support the view that the RT-based classifier characterized the neural signal coding for decision formation.

Applying MVPA to decode unisensory signal cue from noise unveils two processing stages. The later was equivalent to the decision formation stage characterized by the RT-based classifier. The former neural signal was reminiscent of the early target selection signal, typically related to sensory encoding (Luck and Hillyard, 1994; Gamble and Woldorff, 2015; Loughnane et al., 2016). As measured in the present study, this component is also modulated by the amount of sensory evidence provided by the cue and is characterized by a parieto-occipital negativity. This early target selection component has been recently found to modulate the onset and the rate of the CPP (Loughnane et al., 2016) and thereby appears as a processing step mediating decision formation. Altogether, our two decoding analyses of unisensory-cue trials (*i.e.*, against audio-visual noise and RT-based) are concordant and trace the temporal trajectory of neural processes from sensory encoding to decision related signal gradually building-up before the response. Finally, our results show that this chain of processes is similar between modality, and thus generalize findings from human EEG/MEG studies conducted in the visual domain (Ratcliff et al., 2009; Wyart et al., 2012; Mostert et al., 2016).

### Multisensory signal cues accelerate both sensory and decision processes

To portray the dynamic of brain processes following the presentation of multisensory-cues as compared to unisensory-cue, we performed a temporal generalization across conditions. The critical advantage of generalizing in time relies on the fact that unlike classical approach comparing the same time points between conditions, temporal generalization matrix provides comparisons between all time-points allowing to relate brain operations occurring at different latencies. Cross conditions decoding revealed that classifiers trained on unisensory trials were able to decode multisensory trials successfully, it was effective at both processing stages (*i.e.*, sensory encoding and decision formation) and performances were high for the two types of decoding that we conducted (*i.e.*, against audio-visual noise trials and based on response times regression). This cross-condition decoding demonstrated that unisensory and multisensory decision making share the same trajectory. However, time generalization revealed that decoding performance of multisensory trials was more accurate and reached significance at earlier latencies than the ones they were trained on (*i.e.*, unisensory trials). Critically, the speed-up was not limited to a given period but increased with time along the course of processing. Thus, our results demonstrate for the first time that the combination of multisensory-cues accelerate neural processing dynamics during sensory encoding as well as during decision formation (as depicted in the third hypothesis depicted in Figure 1A).

Acceleration during sensory encoding was suggested by a body of work describing how multisensory integration influences early sensory processing (Foxe and Schroeder, 2005; van Wassenhove et al., 2005; Lakatos et al., 2007; Kayser et al., 2008; Romei et al., 2009; Cappe et al., 2012; Mercier et al., 2013, 2015). From there it could be hypothesized that in the case of congruent multisensory source of information, a speed-up at the sensory stage caused by multisensory integration would pass on decision formation (first hypothesis depicted in Figure 1A). Such influence of sensory encoding stage onto decision formation was further supported by a recent study that established a link between the N2 and the CPP: in the context of unpredictable source of information the latency of early target selection signal preludes to faster RTs through earlier evidence accumulation as measured by the CPP (Loughnane et al., 2016). Equally, acceleration during decision formation was suggested by works on perceptual decision making where neural measure of the rate of decision formation varies with the amount of sensory evidence (Heekeren et al., 2008; Romo and de Lafuente, 2013; Hanks and Summerfield, 2017; O’Connell et al., 2018). Also, in the case of sensory evidences originating from multiple modalities, it can be hypothesized that the buildup rate of decision would be quicken: that is decision formation would be shortened by multisensory-cues (second hypothesis depicted in Figure 1A). Our study conciliates these two non-exclusive hypotheses: we show that when cues are available in two modalities sensory encoding is speedup which leads decision formation to happen earlier, next decision formation is further quicken by multisensory evidences fostering the buildup rate of decision formation (third hypothesis depicted in Figure 1A). This acceleration of neural processes during both sensory encoding and decision formation highly suggests that multisensory integration is at play during each processing stage.

### Multisensory integration arises during sensory encoding and decision formation

Multisensory interactions are pervasive in human brain and complete different processes along the cortical hierarchy (Ghazanfar and Schroeder, 2006; Werner and Noppeney, 2010; Rohe and Noppeney, 2015). Sensory regions are the earliest cortical stages of multisensory convergence (Foxe and Schroeder, 2005; Cappe et al., 2009). In these areas, neural activity is modulated by cross-modal inputs (Lakatos et al., 2007; Kayser et al., 2008; Mercier et al., 2013, 2015), modulations which mainly relate to low level features of the different sensory inputs such as their co-occurrence in a short temporal window and/or in a small region of space. Sensory regions also closely interact with higher order areas (*e.g.*, parietal and frontal associative cortex) which mediate integration processes at a higher level, for instance to evaluate the congruency of sensory signals, their reliability or task relevance (Noppeney et al., 2010; Rohe and Noppeney, 2016; Kayser et al., 2017). In our study, to examine neural activity that was specific to the integration of multisensory-cues leading to behavioral benefit, we used a linear classifier to decode multisensory-cues trials from unisensory-cue trials. The neural signature of multisensory integration was found to be concomitant with the previously characterized stages of sensory encoding or decision formation and presented similar topographies, while slightly differing with a right hemisphere dominance. The earliest period of multisensory integration was characterized by a negative parieto-occipital negativity comparable to the early target selection signal (*i.e.*, N2 see above), while the later period of multisensory integration was characterized by a positive centro-parietal positivity alike decision formation signal (*i.e.*, CPP see above). Moreover, the strength of early and late activation patterns were correlated respectively with the strength of sensory encoding and decision formation, suggesting a functional link between these processing steps and multisensory integration. Accordingly, our result substantiates the existence of distinct multisensory processes shaping distinct computational stages and demonstrates for the first time that multisensory integration interplay with decision making, not only sequentially by accelerating sensory processing and thereby quickening the onset of decision formation, but also concurrently by speeding up the rate of decision formation. As such, multisensory integration appears as a crucial factor in perceptual decision making which should be taken into account for building a complete understanding of this multifaceted process.

## Conclusion

In the present study we used multiple MVPA approaches to track the processing stages (*i.e.*, sensory encoding and decision formation) of perceptual decision making following unisensory-cue in a detection task and in a categorization task. From there, we applied cross conditions temporal generalization decoding to multisensory-cues conditions and demonstrated that both sensory encoding and decision formation stages were accelerated in the two tasks. Finally, we characterized multisensory integration and reveal that early and late periods of multisensory integration were tightly linked to sensory encoding and decision formation, respectively. In conclusion our study demonstrates in two tasks and for the first time that multisensory signals foster decision making by speeding-up sensory encoding stages and by increasing the rate of decision formation.

