## Supplemental Figures for "The interplay between multisensory integration and perceptual decision making"

### Figure Supplements

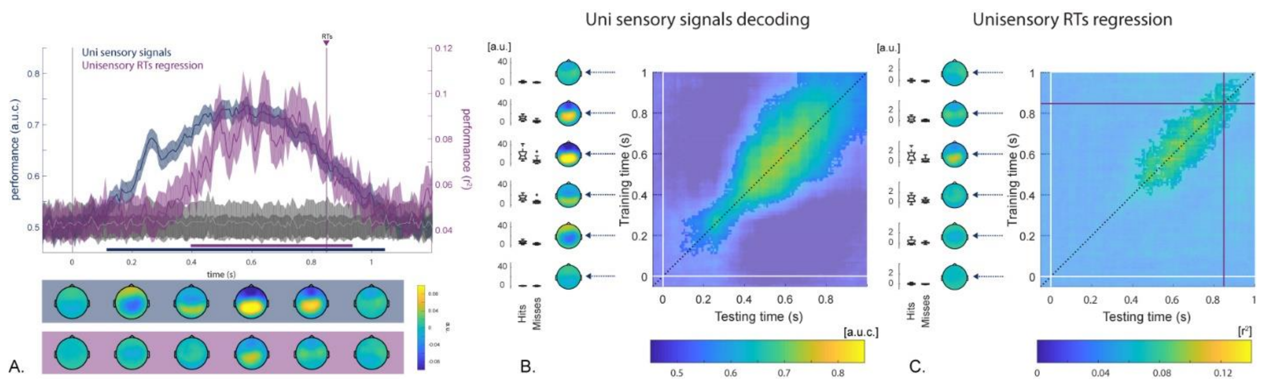


**Supplementary Figure 1: Categorization task, time-resolved pattern classification of unisensory signal encoding and decision formation.**

(A) Mean decoding performance as a function of time (+/- 95% mean confidence intervals). In blue: decoding of unisensory signal cue trials against noise trials. In purple: decoding based on response times. For both, chance levels are depicted in gray. Vertical line represents subject-average response time. Color-coded lines below the x-axis signify statistically significant decoding performance compared to the chance level (corrected for multiple comparisons across time, $p < .005$). Topographical plots underneath depict activation patterns corresponding to the classifier weights (mean over 200 ms time window).

Temporal generalization matrix and corresponding activation patterns depicted for the classifier decoding unisensory trials from trials containing audio-visual noise only (A), and for the classifier decoding response times. Box plots illustrate the consistency of activation patterns across single trials binned by the amount of signal to noise ratio of the cue or between single trials hit or miss.

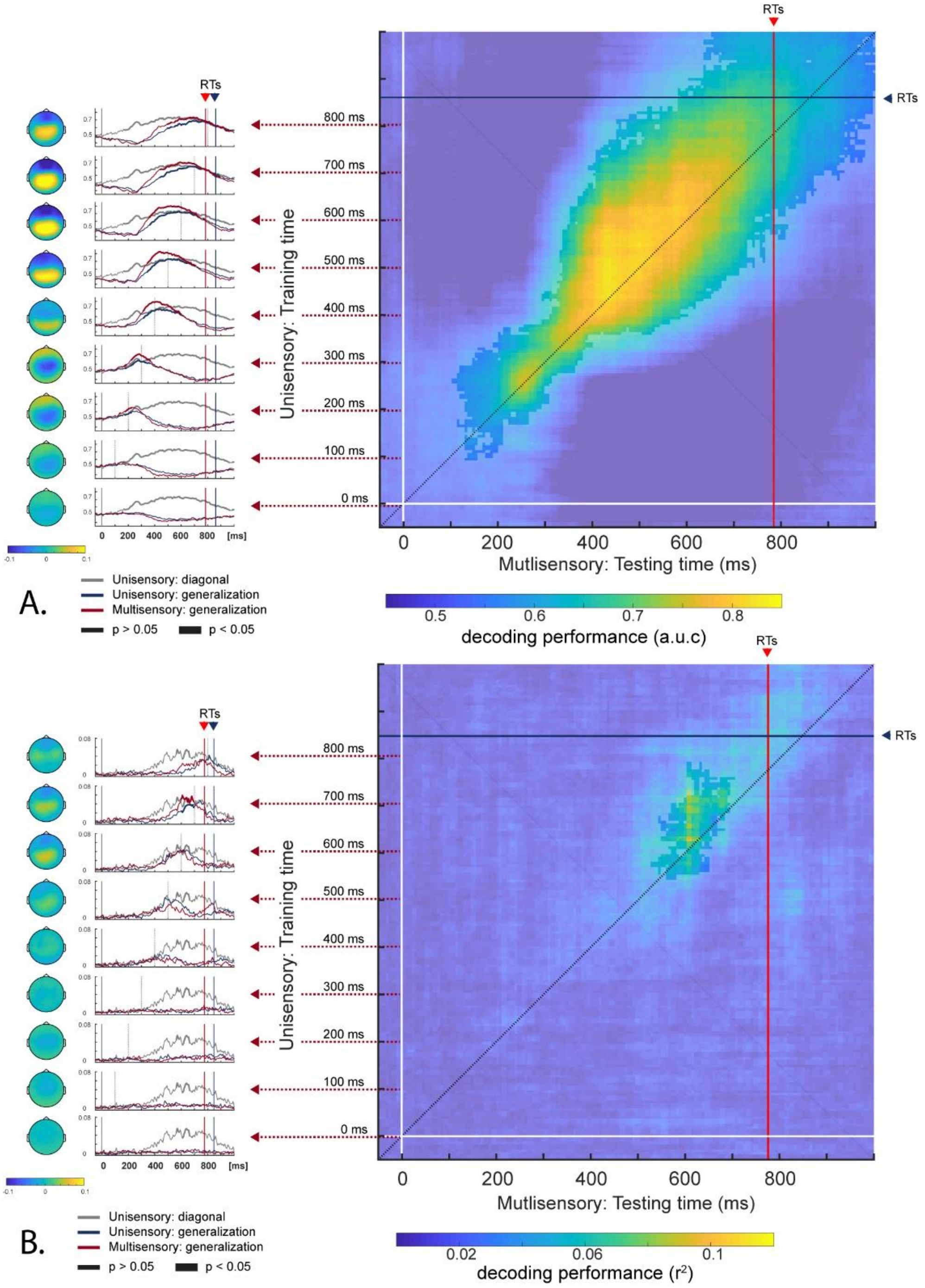


**Supplementary Figure 2: Categorization task, generalization from unisensory to multisensory.** Classifiers trained on unisensory trials were tested (A) to distinguish multisensory signal cue trials vs. trials containing audio-visual noise only, or (B) to decode response times of multisensory-cue trials. Right side: Classifiers trained at a given time (y-axis) are tested at every other time moment (x-axis). Blue horizontal and red vertical lines represent mean response times for unisensory trials and multisensory trials respectively. Left side: Each row represents activation patterns and decoding performance of classifiers tested on multisensory trials (in red) and on unisensory trials (in blue), plotted every 100 ms of training time (with 0 ms on the bottom). The grey lines represent decoding performance for unisensory trials along the diagonal of the temporal generalization matrix (dotted line along the diagonal in supplementary Figure 1).

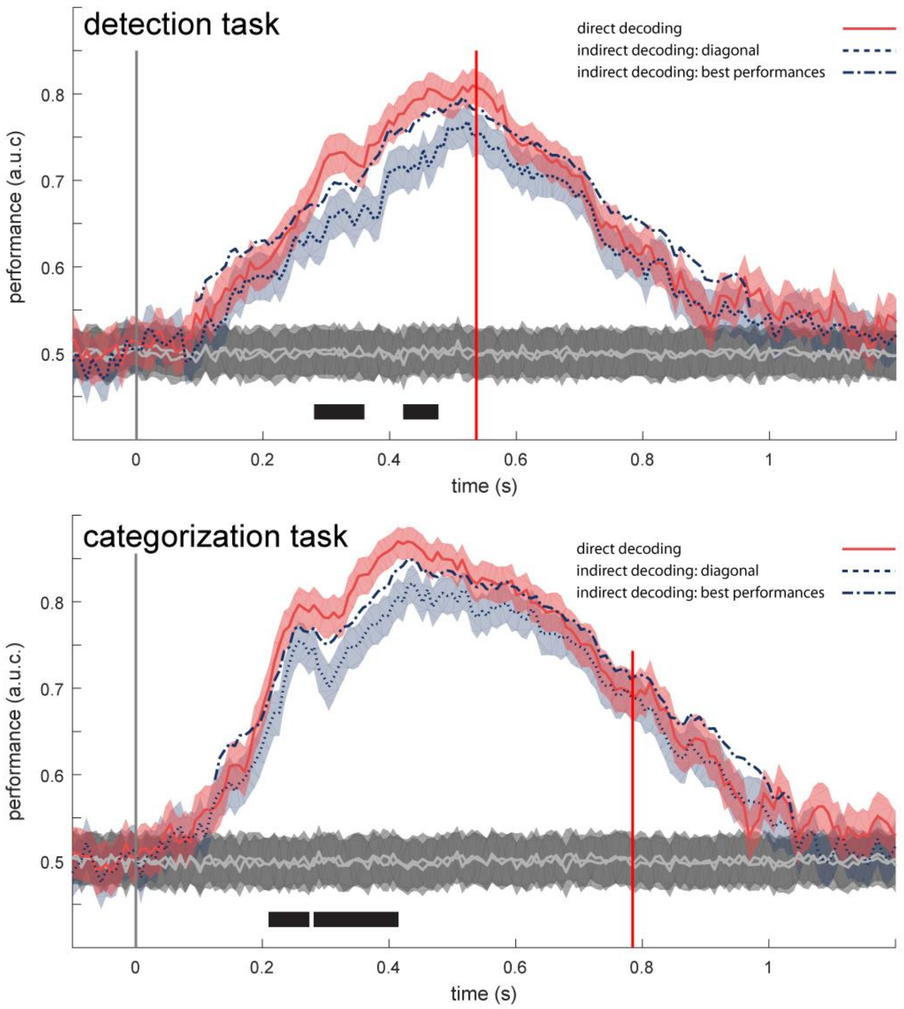


**Supplementary Figure 3: Comparison of direct and indirect decoding performances.**
Time-resolved pattern classification of multisensory signal cues trials versus multisensory noise trials in the detection task (upper panel) and in the categorization task (lower panel). Direct decoding of multisensory signal cues (in red) against multisensory noise is compared to indirect decoding where classifiers were trained on unisensory-cue trial and tested on multisensory-cues trials. Performances of indirect decoding correspond to either the diagonal of the temporal generalization (dotted purple line) of the maximum performances (dashed-dotted purple line). Chance level is represented in gray. Shaded area correspond to mean confidence intervals. Black shapes indicate significant difference between direct decoding and indirect decoding.

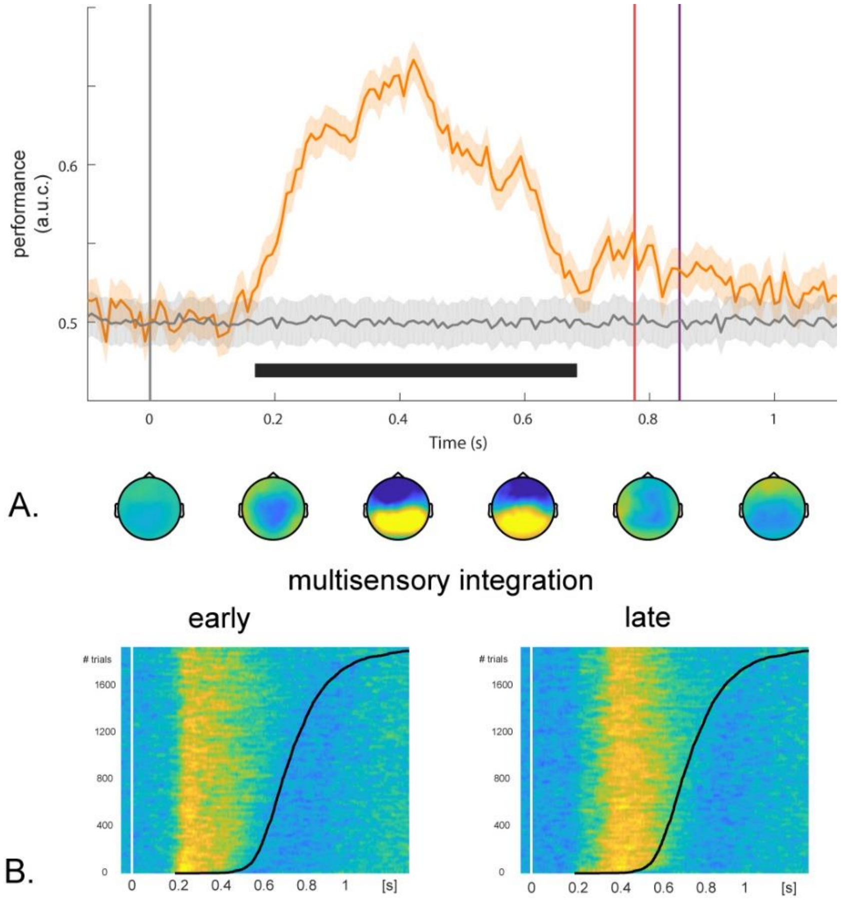


**Supplementary Figure 4: Categorization task, time resolved decoding of multisensory integration.** (A) Mean classification performance (orange line) as a function of time (+/- CI mean; chance levels in gray.), with the black line under the x-axis indicating statistically significant decoding performance compared to the chance level (corrected for multiple comparisons across time, $p < .005$). Topographical maps underneath depict the activation patterns from time-resolved classification procedure (mean over 200 ms time window). Vertical lines represent mean response times for multisensory trials (in red), and unisensory trials (in blue). (B) Single-trial projection of channel weights from classifiers decoding early or late multisensory integration periods. Single trials are sorted by response times (black line).
